# *Bradyrhizobium diazoefficiens* USDA 110-*Glycine max* interactome provides candidate proteins associated with symbiosis

**DOI:** 10.1101/288811

**Authors:** Li Zhang, Jin-Yang Liu, Huan Gu, Yanfang Du, Jian-Fang Zuo, Zhibin Zhang, Menglin Zhang, Pan Li, Jim M. Dunwell, Yangrong Cao, Zuxin Zhang, Yuan-Ming Zhang

**Author notes:** Correspondences or) College of Plant Science and Technology, Huazhong Agricultural University, Wuhan, China. These authors contributed equally to this work (Li Zhang, Jin-Yang Liu). **Data Availability Statement** All the datasets analyzed were from previously published datasets. Supporting Information may be found in additional files.

## Abstract

Although the legume*-*rhizobium symbiosis is a most important biological process, there is a limited knowledge about the protein interaction network between host and symbiont. Using interolog and domain-based approaches, we constructed an inter-species protein interactome with 5115 protein-protein interactions between 2291 *Glycine max* and 290 *Bradyrhizobium diazoefficiens* USDA 110 proteins. The interactome was validated by expression pattern analysis in nodules, GO term semantic similarity, and co-expression analysis. One sub-network was further confirmed using luciferase complementation image assay. In the *G. max-B. diazoefficiens* interactome, bacterial proteins are mainly ion channel and transporters of carbohydrates and cations, while *G. max* proteins are mainly involved in the processes of metabolism, signal transduction, and transport. We also identified the top ten highly interacting proteins (hubs) for each of the two species. KEGG pathway analysis for each hub showed that two 14-3-3 proteins (SGF14g and SGF14k) and five heat shock proteins in *G. max* are possibly involved in symbiosis, and ten hubs in *B. diazoefficiens* may be important symbiotic effectors. Subnetwork analysis showed that 18 symbiosis-related SNARE proteins may play roles in regulating bacterial ion channels, and SGF14g and SGF14k possibly regulate the rhizobium dicarboxylate transport protein DctA. The predicted interactome and symbiosis proteins provide a valuable basis for understanding the molecular mechanism of root nodule symbiosis in soybean.

## Introduction

Rhizobia are gram-negative soil bacteria and have the ability to establish a nitrogen-fixing symbiosis on the roots of legume plants [1, 2]. This legume*-*rhizobium symbiosis is of great agronomic importance and allows the plant to grow successfully in the absence of externally supplied nitrogen fertilizer [1]. Using the legume-rhizobium symbiosis to improve soil fertility is also an effective way to rehabilitate infertile land.

Among rhizobia, *Bradyrhizobium diazoefficiens* USDA 110 (previously named *Bradyrhizobium japonicum* USDA 110) is the most agriculturally important rhizobial bacterium as it is able to specifically infect soybean (*Glycine max*), one of the important legume plants in the world, and form a nitrogen-fixing symbiosis [3]. Furthermore, *G. max-B. diazoefficiens* is one of the most studied soybean*-*rhizobium symbiotic models [4]. Given the importance of such unique feature of legumes, further studies on the mechanisms of the soybean-rhizobium symbiosis are of particular interest. Importantly, the genome sequences of both *B. diazoefficiens* USDA 110 and *G. max* are now available [3, 5], and provide an opportunity to better understand the mechanism of symbiotic features in terms of genomics and proteomics.

In *B. diazoefficiens* USDA 110, several genes related to various stages of the symbiosis process have been identified [3]. In soybean, comparative genomics analysis of legumes also predicted several nodulin genes [5]. Additionally, microarray approaches and RNA-seq analysis in soybean revealed a large number of genes differentially regulated during the symbiosis [4,6,7]. However, none of the above studies have focused on the complex interactions between candidate symbiosis-related genes. Generally, the proteins in the symbiosis process function as a complex network, which combines complex chemical, physical and biological interactions between rhizobial bacteria and their host plants [8]. To better elucidate the complex microbial communities and investigate the mechanism of nitrogen-fixing symbiosis, it is necessary to construct the protein interactions between rhizobium and their host legume plants [9].

For any host-microbe system (including legume-rhizobium symbiosis and host-pathogen system), it is important to understand the mechanism by which the symbiotic or pathogenic bacteria can infect its host. As is known, one of the infection processes of any host-pathogen system is via protein-protein interactions (PPIs) between pathogen proteins and their host proteins [10]. PPIs are the associations of proteins with each other. They play crucial roles in the infection process and in initiating a defense response [11–13]. To date there have been several studies that have focused on the interactions among the protein networks of a host and a pathogen, and identified many new candidate proteins associated with the invasion [11,13–16]. However, PPI network analyses between two species have not been applied to legume-rhizobium symbiosis studies. Therefore, we attempted to construct the PPI interactome between soybean proteins and *B. diazoefficiens* USDA 110 proteins at a genome scale; such an investigation represents a critical step for studying the molecular basis of soybean*-*rhizobium symbiosis.

In the past decade, a series of computational approaches for PPI prediction have been developed [16, 17], and these now play important roles in complementing the various experimental approaches. The existing computational approaches for PPI prediction have exploited diverse data features, which include domain and motif information [18–21], network topology [21, 22], gene ontology (GO) [18–20], gene expression [18, 19], protein sequence similarity [14, 23], and pathway analysis [24]. At present, the interolog and domain-based approaches [25–27] are widely used [14,15,28]. The interolog method is based on protein sequence similarity to conduct the PPI prediction, which maps interactions in the source organism onto the target organism to find possible interactions in the target organism [25, 26]. The domain-based method uses domain interaction information and relies on the principle that if a protein pair contains an interacting domain pair, the two proteins are expected to interact with each other [27].

In this study, we predicted a protein-protein interaction network between *G. max* and *B. diazoefficiens* USDA 110 using both interolog and domain-based methods. GO annotation and gene expression data were utilized to validate the quality of the predicted PPI network. PANTHER overrepresentation test and KEGG pathway enrichment analysis were conducted to determine the biological function of the *B. diazoefficens* and *G. max* proteins predicted in the PPI network. We analyzed the subnetworks of the protein interactome to identify the candidate proteins possibly related to the soybean-rhizobium symbiosis, and used luciferase complementation image (LCI) assay [29, 30] to confirm a subnetwork with two 14-3-3 proteins. In addition, we discuss how these predicted PPIs can help us to better understand this process.

## Results

### Network construction

Based on the well-studied experimental PPIs of seven model organisms: *Arabidopsis thaliana*, *Caenorhabditis elegans*, *Drosophila melanogaster*, *Escherichia coli* K12, *Homo sapiens*, *Mus musculus* and *Saccharomyces cerevisiae*, the PPIs between *G. max* and *B. diazoefficiens* were predicted in this study. To make use of more comprehensive information, we obtained the PPIs of seven organisms from multiple databases: BioGrid [31], DIP [32], HPRD [33], IntAct [34], MINT [31] and TAIR [35]. An ID dictionary was obtained from BioGrid to provide cross-database ID mapping. For mismatching IDs, we corrected manually in the Uniprot ID mapping server. As a result, we incorporated 44702 PPIs with 9948 proteins in *A. thaliana*, 28791 PPIs with 11543 proteins in *C. elegans*, 78383 PPIs with 9438 proteins in *D. melanogaster*, 24460 PPIs with 3358 proteins in *E. coli* K12, 281387 PPIs with 15937 proteins in *H. sapiens*, 31010 PPIs with 8567 proteins in *M. musculus*, and 311333 PPIs with 6149 proteins in *S. cerevisiae* (Table S1).

All the 56044 *G. max* and 8317 *B. diazoefficiens* USDA 110 proteins were used to conduct a genome-wide PPI prediction. Among 8317 *B. diazoefficiens* proteins, 2356 proteins are secreted or membrane proteins (Table S2), which have the possibility to interact with *G. max* proteins. Using the pipeline shown in Figure 1 and filtered by above 2356 secreted or membrane proteins, 5115 PPIs between 2291 soybean proteins and 290 *B. diazoefficiens* USDA 110 proteins were predicted (Figure S1; Table S3). In addition, 233545 intra-species PPIs in soybean (Table S4) and 11106 intra-species PPIs in *B. diazoefficiens* USDA 110 (Table S5) were predicted. In summary, there were a total of 249766 PPIs, including inter- and intra-species PPIs, and 54471 PPIs (21.81%) were found in more than one species or experiment. All predicted interactions and the detailed annotation information of the proteins are available in Tables S3 to S5.

**Figure 1.**
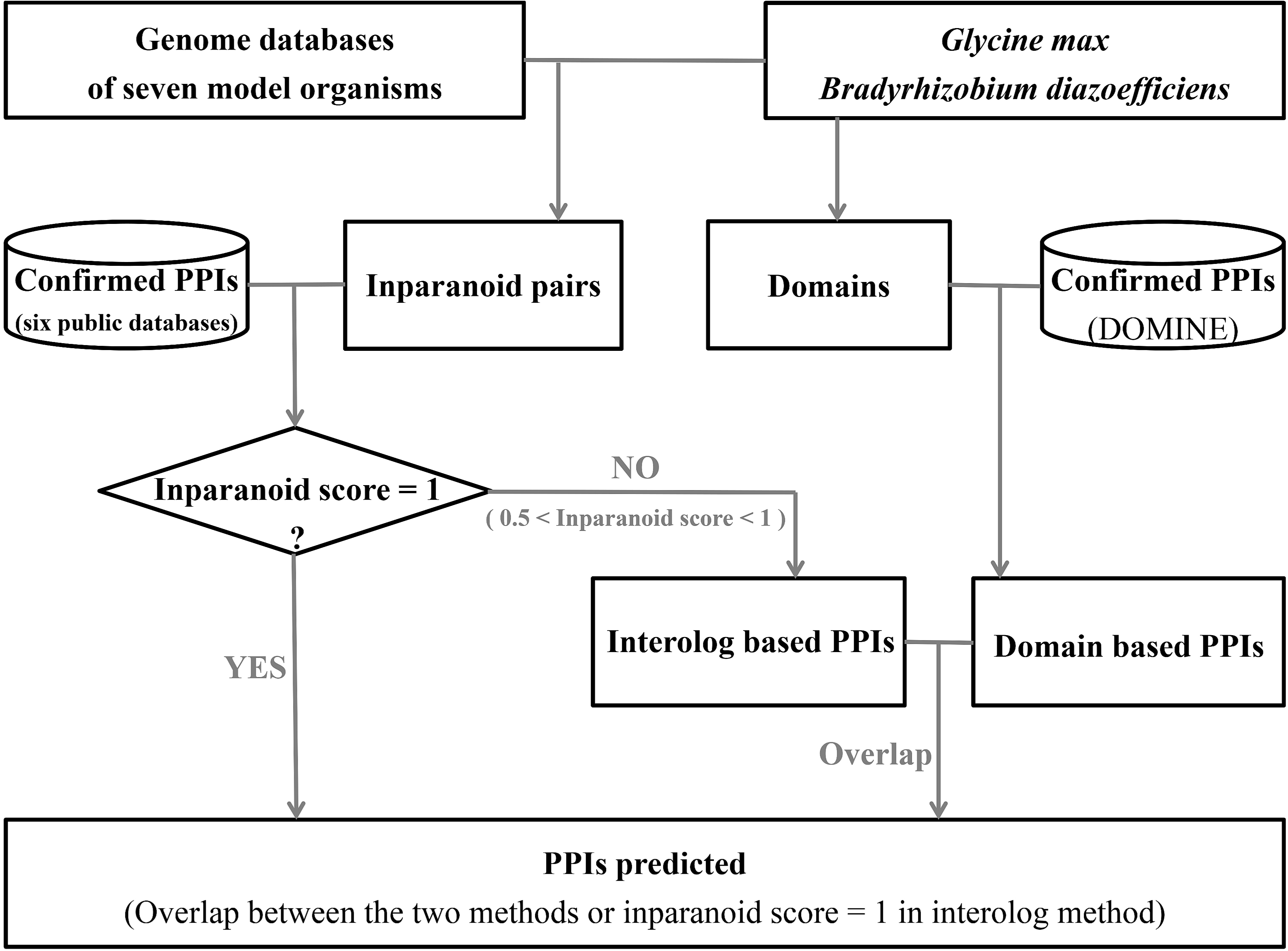
The prediction pipeline of the protein-protein interaction networks.

### Quality assessment of protein–protein interactions

To date, few experimental PPIs between *B. diazoefficiens* USDA 110 and *G. max* have been identified, so it is difficult to validate the predicted PPI network by experimental approaches. For this reason, computational biology approaches were used to validate the quality of the predicted PPI network. In this study, we analyzed the gene expression pattern in nodules of all the soybean and rhizobium proteins in the *G. max-B. diazoefficiens* interactome. Furthermore, we conducted GO term semantic similarity [23, 36] and co-expression analysis [28, 37] of the intra-species PPI interactome. The results were used to deduce the quality of the *G. max*-*B. diazoefficiens* interactome, owing to the same methodologies.

### Expression pattern in nodules

The interaction between rhizobium and its host legume results in the formation of a novel plant organ, the nodule. In nodules, the legume host interacts with rhizobium and exchanges photosynthetic products for ammonia from the rhizobial bacteria [38]. Thus, the predicted 5115 interactions between *B. diazoefficiens* and *G. max* are more likely to occur in nodules. In other words, most genes that encode the 2291 soybean proteins and the 290 *B. diazoefficiens* proteins in 5115 PPIs should be expressed in nodules. Analysis of the transcriptome data showed that 71.80% (1644) soybean genes were expressed in nodules with FPKM > 5. However, for the whole genome, the percent of genes expressed in nodules with FPKM > 5 is only 33.34% (18686 genes of the entire genome, which has 56045 genes). This indicates that most soybean genes in the above predicted network were indeed significantly expressed in nodules.

In previous studies, genome-wide analysis of *B. diazoefficiens* genes in symbiosis bacteroids was conducted at the transcriptome [39, 40] and protein [41] levels. And these datasets were also used to investigate the expression patterns of 290 *B. diazoefficiens* genes in soybean root nodules. As a result, 172 (59.31%) genes were found to be expressed in symbiosis bacteroids (Table S6).

### Functional similarity based on GO annotation

Two interacting proteins would have similar or related functions and should share some common GO annotations [23,28,36]. Thus, GO annotation information of two interacting proteins was used to measure the accuracy of our prediction. Among 56044 soybean genes, 30023 (53.57%) genes were annotated with at least one GO term in any of the three GO categories (molecular function, biological process, and cellular component). Of all the 233545 soybean PPIs, 128862, 66369 and 26135 PPIs were annotated in the categories of molecular function, biological process, and cellular component (125086, 63581, 25007 non-self interactions), respectively (Table S4).

To measure the semantic similarity between GO terms and to evaluate the reliability of predicted PPIs, three functional similarity scores, sim_JC_^BP^, sim_JC_^MF^ and sim_JC_^CC^, were calculated using non-self interactions in each GO category. Meanwhile, randomly selected protein pairs of the same size served as a control. As a result, significant differences for each of three simJC scores between predicted PPIs and randomly selected protein pairs were observed (Figure 2). All the proportions of score 1.0 in sim_JC_^BP^, sim_JC_^MF^, and sim_JC_^CC^ were significantly higher in predicted soybean PPIs than those in randomly selected protein pairs, indicating that the predicted interaction network indeed preferentially connects functionally related proteins.

**Figure 2.**
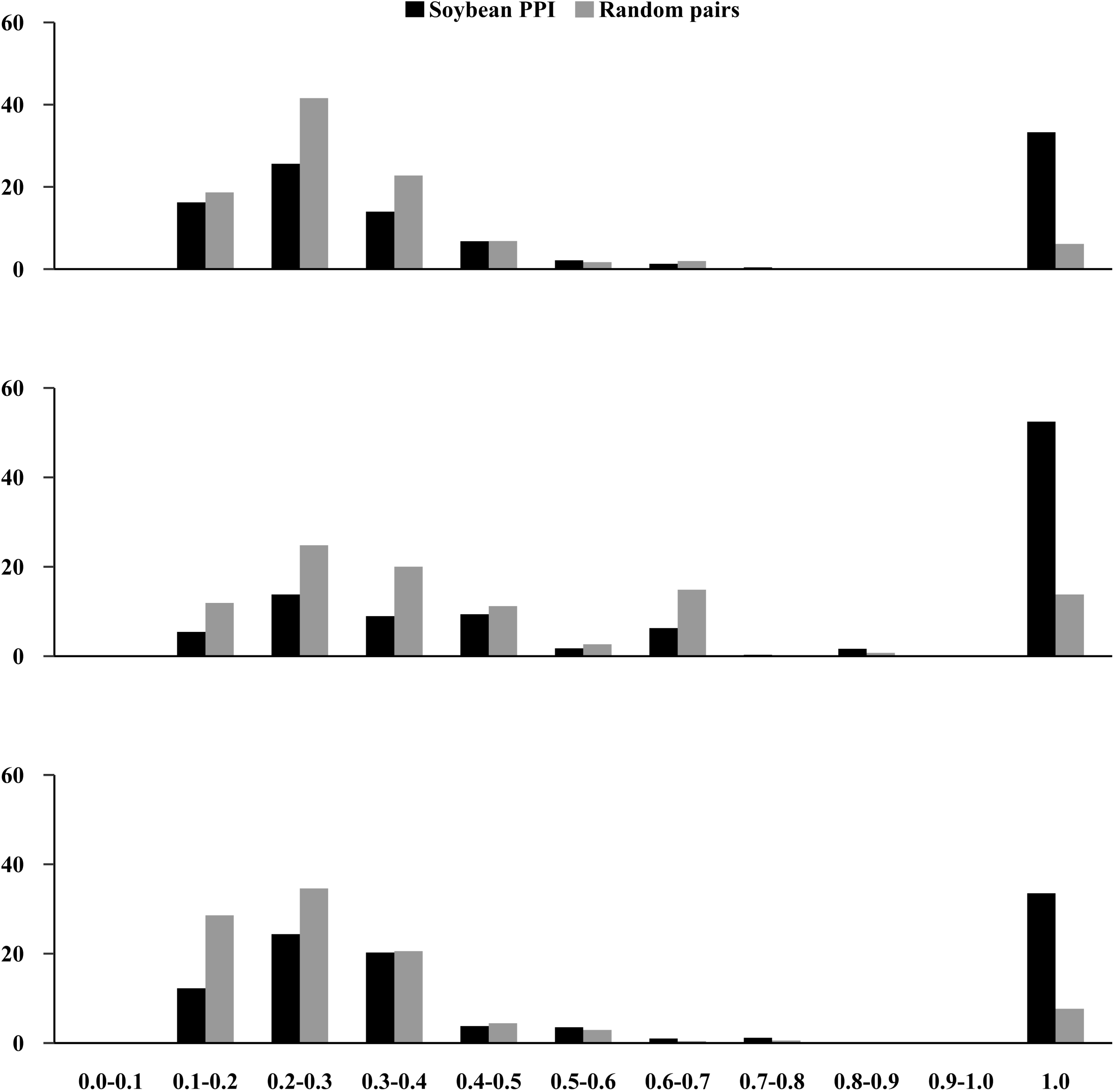
Distribution of semantic similarity scores between GO terms of two proteins: sim_JC_^BP^**, sim_JC_^MF^ and sim_JC_^CC^**. A: distribution of sim_JC_^BP^; B: distribution of sim_JC_^CC^; C: distribution of sim_JC_^MF^. Box in black represents predicted protein-protein interactions in soybean; grey box denotes random protein pairs in the soybean genome.

### Co-expression of predicted soybean PPIs

Levels of mRNA expression have some relationship with protein-protein interactions [42]. The interacting proteins tend to have correlated gene expression patterns, especially for subunits of the same protein complex [28,37,43]. Thus, we investigated the relationship of our predicted intra-species PPIs with mRNA expression levels in soybean. In this study, we used the transcriptome data from nine tissues of *G. max* to investigate expression correlation between two interacting proteins. The co-expression level of two interacting proteins was calculated by a widely used measure, the Pearson correlation coefficient (PCC) [44].

Among 233545 soybean intra-species PPIs (Table S3), 216097 PCC scores were successfully calculated. Among these scores, 23.84% (51524) protein interactions had a high PCC score (*r* > 0.6). In randomly selected protein pairs, however, the proportion was only 13.80%. This implies that the predicted interacting pairs have a significant co-relationship and the predicted PPI networks have high reliability. For conserved PPIs identified from more than one species or experiment, 34.72% had a high PCC score (*r* > 0.6), indicating a higher reliability. This is consistent with the conclusion that protein interactions detected by more than one high-throughput interaction assay are more accurate [36, 45].

### Conserved PPIs identified in more than two species

Common protein interactions predicted from multiple species can be considered as evolutionarily conserved interactions that have very high confidence [36]. In this study, we detected common protein interactions from more than two species. As a result, 60 conserved PPIs including 54 *G. max* proteins and 21 *B. diazoefficiens* proteins in *G. max-B. diazoefficiens* interactome were found (Figure S2). Among these 54 *G. max* proteins, more importantly, 49 proteins were expressed with FPKM > 5 in the underground tissues (root, root hair and nodule) and 24 proteins had high expression levels with FPKM > 100 in the underground tissues.

### Function enrichment analysis of proteins in *G. max-B. diazoefficiens* interactome

To determine whether any biological function biases exist in the *B. diazoefficens* and *G. max* proteins in the predicted PPI network, we classified the proteins using the PANTHER overrepresentation test and conducted KEGG pathway enrichment analysis. The corresponding results with Bonferroni correction are listed in Tables 1 and 2, respectively.

**Table 1.**
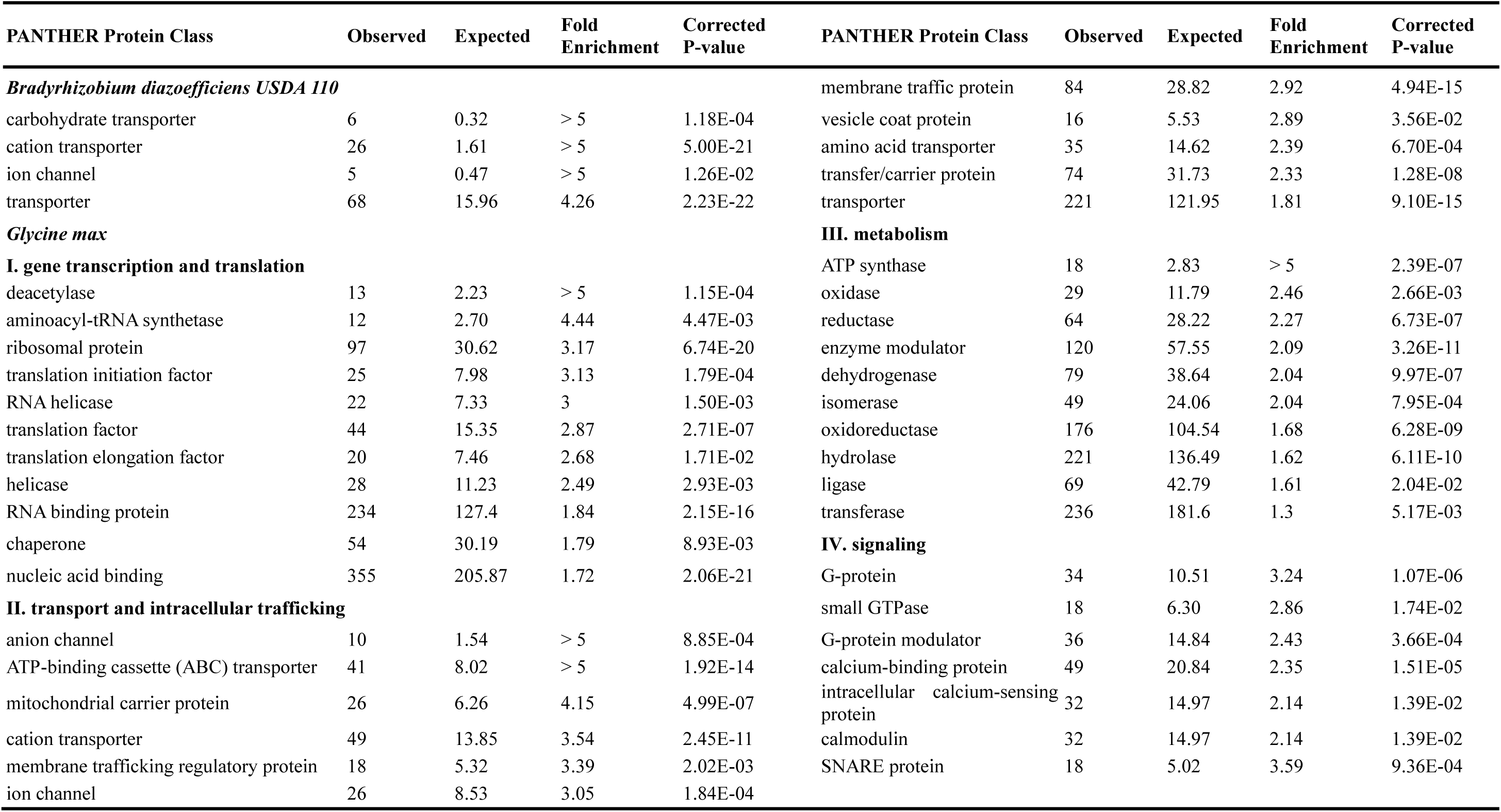
Classification of proteins in predicted PPIs between soybean and B. diazoefficiens USDA 110 by PANTHER overrepresentation test.

| PANTHER Protein Class | Observed | Expected | Fold Enrichment | Corrected P-value | PANTHER Protein Class | Observed | Expected | Fold Enrichment | Corrected P-value |
| --- | --- | --- | --- | --- | --- | --- | --- | --- | --- |
| *Bradyrhizobium diazoefficiens* USDA 110 |  |  |  |  | membrane traffic protein | 84 | 28.82 | 2.92 | 4.94E-15 |
| carbohydrate transporter | 6 | 0.32 | > 5 | 1.18E-04 | vesicle coat protein | 16 | 5.53 | 2.89 | 3.56E-02 |
| cation transporter | 26 | 1.61 | > 5 | 5.00E-21 | amino acid transporter | 35 | 14.62 | 2.39 | 6.70E-04 |
| ion channel | 5 | 0.47 | > 5 | 1.26E-02 | transfer/carrier protein | 74 | 31.73 | 2.33 | 1.28E-08 |
| transporter | 68 | 15.96 | 4.26 | 2.23E-22 | transporter | 221 | 121.95 | 1.81 | 9.10E-15 |
| *Glycine max* |  |  |  |  | **III. metabolism** |  |  |  |  |
| **I. gene transcription and translation** |  |  |  |  | ATP synthase | 18 | 2.83 | > 5 | 2.39E-07 |
| deacetylase | 13 | 2.23 | > 5 | 1.15E-04 | oxidase | 29 | 11.79 | 2.46 | 2.66E-03 |
| aminoacyl-tRNA synthetase | 12 | 2.70 | 4.44 | 4.47E-03 | reductase | 64 | 28.22 | 2.27 | 6.73E-07 |
| ribosomal protein | 97 | 30.62 | 3.17 | 6.74E-20 | enzyme modulator | 120 | 57.55 | 2.09 | 3.26E-11 |
| translation initiation factor | 25 | 7.98 | 3.13 | 1.79E-04 | dehydrogenase | 79 | 38.64 | 2.04 | 9.97E-07 |
| RNA helicase | 22 | 7.33 | 3 | 1.50E-03 | isomerase | 49 | 24.06 | 2.04 | 7.95E-04 |
| translation factor | 44 | 15.35 | 2.87 | 2.71E-07 | oxidoreductase | 176 | 104.54 | 1.68 | 6.28E-09 |
| translation elongation factor | 20 | 7.46 | 2.68 | 1.71E-02 | hydrolase | 221 | 136.49 | 1.62 | 6.11E-10 |
| helicase | 28 | 11.23 | 2.49 | 2.93E-03 | ligase | 69 | 42.79 | 1.61 | 2.04E-02 |
| RNA binding protein | 234 | 127.4 | 1.84 | 2.15E-16 | transferase | 236 | 181.6 | 1.3 | 5.17E-03 |
| chaperone | 54 | 30.19 | 1.79 | 8.93E-03 | **IV. signaling** |  |  |  |  |
| nucleic acid binding | 355 | 205.87 | 1.72 | 2.06E-21 | G-protein | 34 | 10.51 | 3.24 | 1.07E-06 |
| **II. transport and intracellular trafficking** |  |  |  |  | small GTPase | 18 | 6.30 | 2.86 | 1.74E-02 |
| anion channel | 10 | 1.54 | > 5 | 8.85E-04 | G-protein modulator | 36 | 14.84 | 2.43 | 3.66E-04 |
| ATP-binding cassette (ABC) transporter | 41 | 8.02 | > 5 | 1.92E-14 | calcium-binding protein | 49 | 20.84 | 2.35 | 1.51E-05 |
| mitochondrial carrier protein | 26 | 6.26 | 4.15 | 4.99E-07 | intracellular calcium-sensing protein | 32 | 14.97 | 2.14 | 1.39E-02 |
| cation transporter | 49 | 13.85 | 3.54 | 2.45E-11 | calmodulin | 32 | 14.97 | 2.14 | 1.39E-02 |
| membrane trafficking regulatory protein | 18 | 5.32 | 3.39 | 2.02E-03 | SNARE protein | 18 | 5.02 | 3.59 | 9.36E-04 |
| ion channel | 26 | 8.53 | 3.05 | 1.84E-04 |  |  |  |  |  |

**Table 2.** KEGG pathway enrichment analysis of proteins in PPIs between soybean and *B. diazoefficiens* USDA 110.

| *Glycine max* |  |  |  |  | *Bradyrhizobium diazoefficiens* USDA 110 |  |  |  |  |
| --- | --- | --- | --- | --- | --- | --- | --- | --- | --- |
| KEGG Term | KEGG ID | Input number | Background number | Corrected P-Value | KEGG Term | KEGG ID | Input number | Background number | Corrected P-Value |
| Oxidative phosphorylation | gmx00190 | 72 | 237 | 9.35E-09 | Oxidative phosphorylation | bja00190 | 20 | 66 | 1.46E-10 |
| Phagosome | gmx04145 | 52 | 165 | 7.00E-07 | Protein export | bja03060 | 10 | 20 | 5.58E-07 |
| Protein export | gmx03060 | 36 | 91 | 8.57E-07 | Two-component system | bja02020 | 20 | 168 | 5.79E-05 |
| Protein processing in endoplasmic reticulum | gmx04141 | 81 | 375 | 6.27E-05 | Peptidoglycan biosynthesis | bja00550 | 6 | 24 | 0.003838 |
| N-Glycan biosynthesis | gmx00510 | 27 | 74 | 9.75E-05 | Glycerophospholipid metabolism | bja00564 | 6 | 24 | 0.003838 |
| Ribosome | gmx03010 | 109 | 595 | 0.000594 | ABC transporters | bja02010 | 22 | 299 | 0.007615 |
| Biosynthesis of amino acids | gmx01230 | 77 | 429 | 0.011763 | Bacterial secretion system | bja03070 | 7 | 44 | 0.009047 |
| Citrate cycle (TCA cycle) | gmx00020 | 26 | 104 | 0.015133 | beta-Lactam resistance | bja01501 | 5 | 21 | 0.009047 |
| Carbon metabolism | gmx01200 | 84 | 488 | 0.016097 |  |  |  |  |  |
| Proteasome | gmx03050 | 24 | 105 | 0.049615 |  |  |  |  |  |

Notes: Input genes and their detailed annotations were available in Supplementary Table S6

### *B. diazoefficens* USDA 110 proteins

In the predicted PPI network, *B. diazoefficens* proteins are mainly ion channel and transporters of carbohydrates and cations (Table 1). As the legume-rhizobium interaction involves the bacterial fixation of atmospheric nitrogen in exchange for plant-produced carbohydrates and all the essential nutrients required for bacterial metabolism [38,46,47], these transporters may provide the opportunities for rhizobial nodulation. KEGG pathway enrichment analysis further showed that bacterial proteins in *G. max-B. diazoefficiens* interactome were involved in pathways associated with symbiosis, such as protein export, peptidoglycan biosynthesis, ABC transporters and the bacterial secretion system (Tables 2 and S7), which are consistent with those in previous studies [48–51].

### *G. max* proteins

Protein classification in soybean showed that proteins interacting with *B. diazoefficiens* were mainly involved in the processes of gene transcription and translation, transport, metabolism, and signal transduction (Table 1). In transport, they were ion channels, ATP-binding cassette (ABC) transporters, mitochondrial carrier proteins and amino acid transporters. In signal transduction, 34 G-proteins, 18 small GTPase, 32 calmodulin and 18 SNARE proteins were present in the predicted PPIs and directly interacted with bacteria (Tables 1 and S8). Moreover, KEGG pathway enrichment analysis showed that soybean proteins in the predicted PPIs were involved in carbon metabolism, tricarboxylic acid cycle and N-glycan biosynthesis (Tables 2 and S7). Consistent with the above observations, Carvalho *et al*. [7] demonstrated that soybean genes involved in signal transduction, transcriptional regulation and primary metabolism were induced by the presence of the rhizobial bacteria. Additionally, by comparing with the *G. max* nodulation-related genes or searching for homologs of *M. truncatula* and *L. japonicus* nodulation-related genes in previous studies [4,5,52], we investigated whether some *G. max* proteins in predicted PPIs are experimentally nodulation-related genes. As a result, 9 soybean nodulation-related genes were identified and their PPIs are list in Table S9. These results suggest that soybean proteins interacting with the rhizobium were involved in various specific areas of metabolism, and the predicted interactions may provide useful information to understand the molecular mechanism of the legume-rhizobium symbiosis.

### Hubs in *G. max-B. diazoefficiens* interactome

In protein–protein interaction networks, most proteins (nodes) connect with few proteins, whereas, a small percentage of proteins interact with a large number of other proteins [53, 54]. Such proteins (nodes) with a large number of interactions are called hubs, and are more essential than proteins with only a small number of interactions. These proteins are known to perform vital roles in various cellular processes under a range of conditions including those caused by host-pathogen interactions [53–56]. In the present study, we listed the top ten hubs of each species in the *G. max-B. diazoefficiens* interactome (Table 3). To further understand the functions of the twenty hubs, we performed KEGG pathway enrichment analysis for the proteins interacting with each of the twenty hubs. These results are listed in Supplementary Table S10.

**Table 3.**
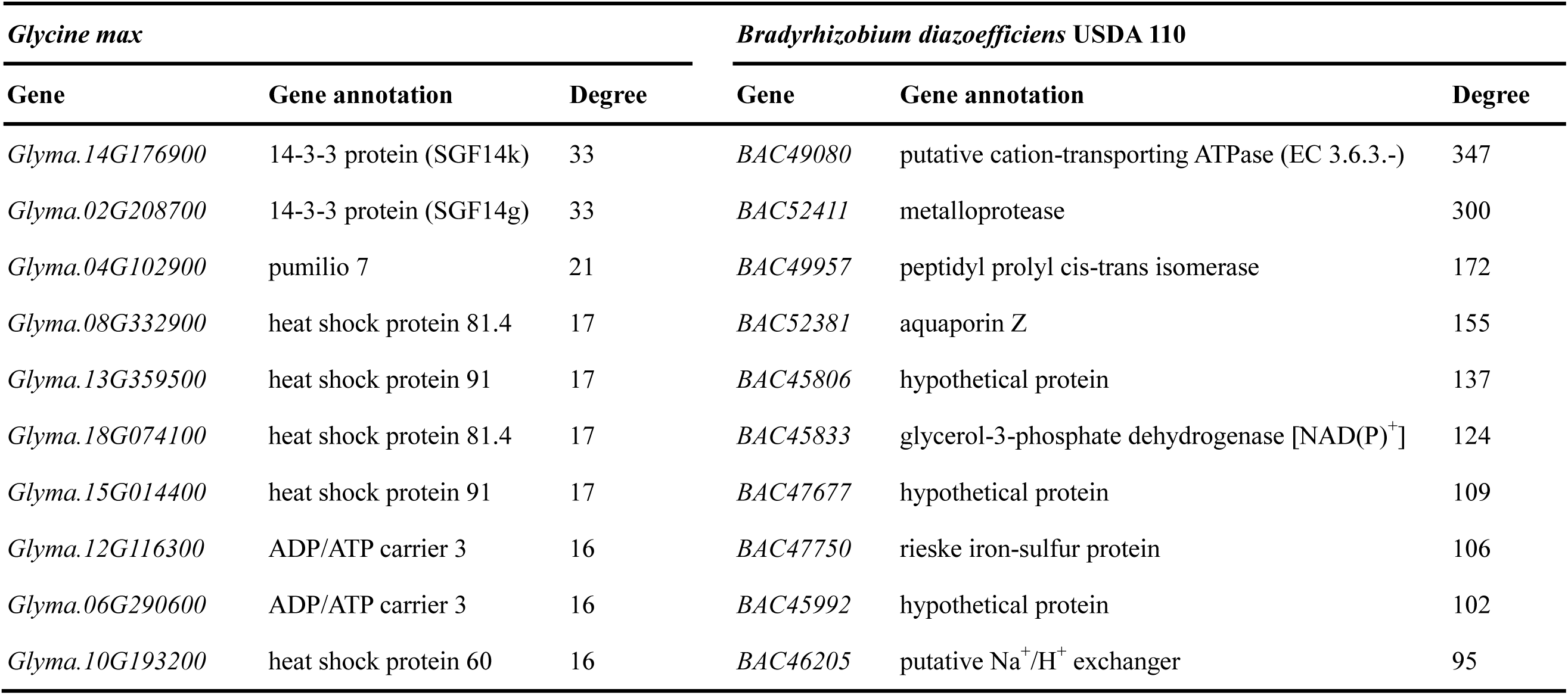
Top ten hubs of *G. max* and *B. diazoefficiens* USDA 110 in the predicted PPI network.

| *Glycine max* |  |  | *Bradyrhizobium diazoefficiens* USDA 110 |  |  |
| --- | --- | --- | --- | --- | --- |
| Gene | Gene annotation | Degree | Gene | Gene annotation | Degree |
| *Glyma.14G176900* | 14-3-3 protein (SGF14k) | 33 | *BAC49080* | putative cation-transporting ATPase (EC 3.6.3.-) | 347 |
| *Glyma.02G208700* | 14-3-3 protein (SGF14g) | 33 | *BAC52411* | metalloprotease | 300 |
| *Glyma.04G102900* | pumilio 7 | 21 | *BAC49957* | peptidyl prolyl cis-trans isomerase | 172 |
| *Glyma.08G332900* | heat shock protein 81.4 | 17 | *BAC52381* | aquaporin Z | 155 |
| *Glyma.13G359500* | heat shock protein 91 | 17 | *BAC45806* | hypothetical protein | 137 |
| *Glyma.18G074100* | heat shock protein 81.4 | 17 | *BAC45833* | glycerol-3-phosphate dehydrogenase [NAD(P)<sup>+</sup>] | 124 |
| *Glyma.15G014400* | heat shock protein 91 | 17 | *BAC47677* | hypothetical protein | 109 |
| *Glyma.12G116300* | ADP/ATP carrier 3 | 16 | *BAC47750* | rieske iron-sulfur protein | 106 |
| *Glyma.06G290600* | ADP/ATP carrier 3 | 16 | *BAC45992* | hypothetical protein | 102 |
| *Glyma.10G193200* | heat shock protein 60 | 16 | *BAC46205* | putative Na<sup>+</sup>/H<sup>+</sup> exchanger | 95 |

In soybean, the top ten hubs included two 14-3-3 proteins, a Pumilio 7 protein, five heat shock proteins (HSPs) and two ADP/ATP carrier proteins (Table 3). The KEGG pathways for the two 14-3-3 proteins contained two-component systems (TCSs), Tryptophan metabolism and Oxidative phosphorylation (Table S10). Pumilio 7 protein and two ADP/ATP carrier proteins were both involved in the processes of Oxidative phosphorylation and Glycerophospholipid metabolism. Three of the five HSPs were enriched to show interaction with bacterial proteins in the metabolism of glycerophospholipids (Table S10), which are important components of membrane lipids in bacteria.

In *B. diazoefficiens*, the ten hubs included BAC49080, BAC52411, BAC49957, BAC52381, BAC45806, BAC45833, BAC47677, BAC47750, BAC45992 and BAC46205 (Table 3). KEGG pathway enrichment analysis showed that seven hubs were involved in carbohydrate metabolism, including N-Glycan biosynthesis, Pyruvate metabolism, Glycolysis and Citrate cycle (Table S10).

### Subnetworks related to symbiosis

Based on an analysis of the PPI networks, we can better understand the web of interactions that takes place inside a cell. One method to better understand the entire network is to partition it into a series of subnetworks. In the present study, we selected two subnetworks that separately contain SNAREs and 14-3-3 proteins for further analysis to identify candidate proteins related to symbiosis (Figure 3).

**Figure 3.**
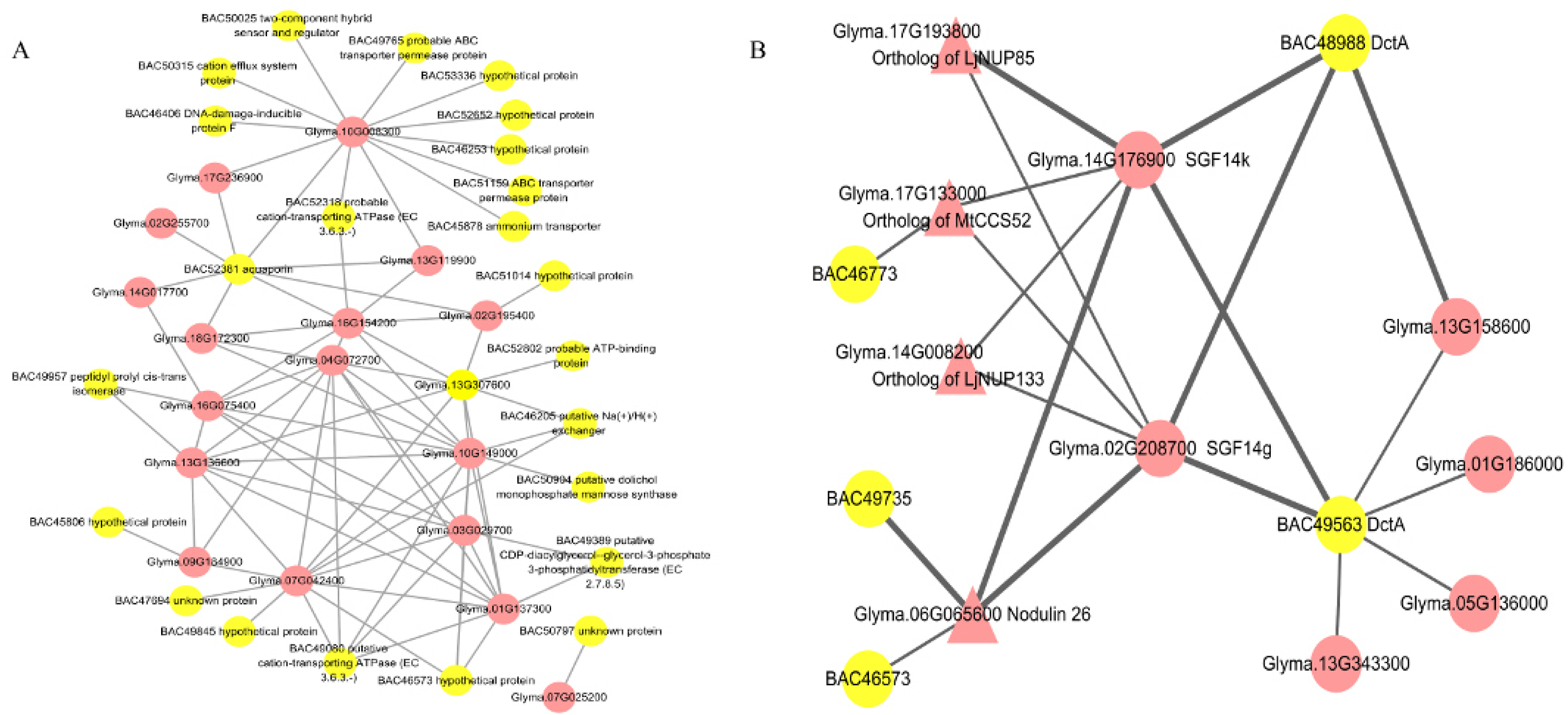
Two PPI sub-networks between soybean (red) and *B. diazoefficiens* USDA 110 (yellow) proteins. A: PPI sub-network between 18 soybean SNARE proteins and *B. diazoefficiens* USDA 110 proteins. B: PPI sub-network of DctA and 14-3-3 proteins. Triangles represent nodulin in soybean. The PPI interactions with bold edges were validated by the LCI assay.

### SNARE proteins

SNARE proteins are vital for signal transduction and membrane fusion in plants [57, 58]. There is now growing evidence that these proteins play crucial roles in symbiosis in legume nodules, such as those in *L. japonicus* [58] and *M. truncatula* [59, 60]. In the present study, 18 SNARE proteins in *G. max* were involved in the predicted *G. max-B. diazoefficens* interactome and closely interacted with *B. diazoefficens* proteins (Figure 3A), suggesting the critical roles of SNAREs in soybean root nodule symbiosis (RNS). Meanwhile, soybean SNAREs interacted with each other in Figure 3A, which was consistent with the results in previous structural studies that SNAREs could form complexes by interacting with other SNAREs [57, 58].

### 14-3-3 protein

14-3-3 proteins are abundant proteins in plants, and are involved in signaling pathways to regulate plant development and response to stimulus. Li and Dhaubhadel [61] identified 18 genes (SGF 14a-r) coding 14-3-3 proteins in the whole soybean genome. Previous studies revealed that two of them (SGF14c and SGF14l) play critical roles in RNS [62] and homologs of SGF14b in *L. japonicus* were located in the peribacteroid membrane [63]. In our study, we found another two 14-3-3 proteins, Glyma.14G176900 (SGF14k) and Glyma.02G208700 (SGF14g), which are hubs that interacted with *B. diazoefficiens* to a high degree (Table 3). More importantly, we found that SGF14k and SGF14g were connected with four soybean nodulation genes, *Glyma.06G065600* (Nodulin26) [64], *Glyma.17G13300* (WD40 protein; homologs of MtCCS52) [65], *Glyma.17G193800* (nucleoporin; homologs of LjNUP85) [66] and *Glyma.14G008200* (nucleoporin; homologs of LjNUP133) (Figure 3B) [67]. The results of the predicted PPIs of SGF14k and SGF14g demonstrated that SGF14k and SGF14g were involved in RNS.

### Validation of a subnetwork containing two 14-3-3 proteins using luciferase complementation image (LCI) assay experiment

Luciferase complementation image (LCI) assay is a well-established method to verify the predicted PPIs in a laboratory setting. To validate the accuracy of the predicted interactions, a subnetwork in Figure 3B was selected to test the interactions *in vivo*. As a result, nine were confirmed (Figure 4). For example, SGF14k was interacted with BAC48988, similarly, SGF14k and BAC49563, SGF14k and Nodulin26, SGF14g and BAC48988, SGF14g and BAC49563, SGF14g and Nodulin26, SGF14g and NUP85, Nodulin26 and BAC49735, and Glyma13G158600 and BAC49563 (Figures 3B and 4). Among the nine pairs of PPIs, six interacting protein pairs are produced between *G. max* and *B. diazoefficens*. More importantly, the interaction between Glyma.06G065600 (Nodulin) and Glyma.17G193800 (nucleoporin; homologs of LjNUP85) has been found to be involved in RNS [64, 66]. Meanwhile, soybean SGF14g and SGF14k were both verified by LCI assay to interact with soybean Nodulin26 and *B. diazoefficiens* proteins BAC48988 and BAC49563, suggesting the critical roles of SGF14g and SGF14k in the establishment of RNS. Additionally, Nodulin26 was also found to be interacted with *B. diazoefficens* protein BAC49735. Taken together, the results demonstrated the reliability of our predicted PPIs, which can provide a useful guideline for future research.

**Figure 4.**
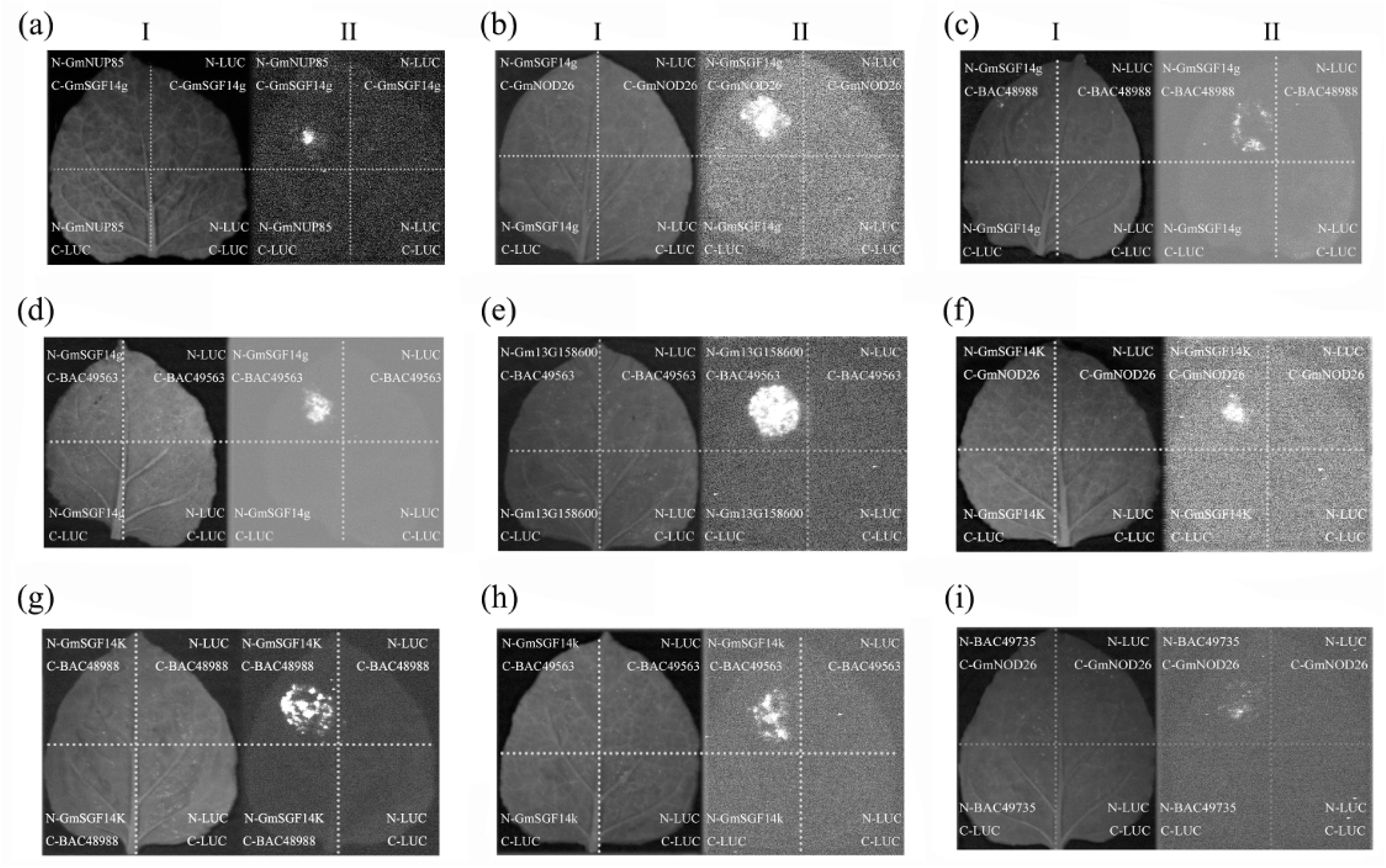
Luciferase complementation image assay of a subnetwork containing two 14-3-3 proteins in *Agrobacterium*-infiltrated *N. benthamiana* leaves under bright field (I) and dark (II) illumination. The C-terminal half and the N-terminal half of LUC were fused to N-gene, N-LUC, C-gene and C-LUC. In (c), the treatment was N-GmSGF14g + C-BAC48988, and the controls were N-LUC + C-BAC48988, N-GmSGF14g + C-LUC, and N-LUC + C-LUC. LUC fluorescence was detected by confocal microscope in *N. benthamiana* fresh leaves. The experiment was repeated three times with similar results. The situation was similar in the others.

## Discussion

### Network validation for the predicted PPIs in this study

The network predicted in this study is relatively reliable. The reasons are as follows. First, nine predicted PPIs in a sub-network containing two 14-3-3 proteins (SGF14g and SGF14k) showed an interaction signal via the LCI assay (Figure 4). Meanwhile, nine soybean nodulation-related genes predicted in this study have been experimentally confirmed to be involved in RNS (Table S9). Additionally, three computational biology approaches were used to validate the predicted network in this study. For example, significantly higher proportions of score 1.0 for the three simJC indicators in predicted soybean PPIs than those in randomly selected protein pairs indicates high quality of the *G. max-B. diazoefficiens* interactome (Figure 2); a significant higher proportion of predicted interaction pairs showed a co-relationship in their gene expression levels (PCC score > 0.6) than did randomly selected protein pairs; soybean genes expressed in nodules with FPKM > 5 had a significantly higher proportion (71.80%) in the predicted network than those (33.34%) in the entire genome, while 59.31% *B. diazoefficiens* genes were found to be expressed in symbiosis bacteroids.

### Soybean proteins in the predicted PPIs were involved with pathways associated with symbiosis

The infection transcriptome analysis confirmed that proteins involved in various areas of metabolism were triggered in the host plant by the presence of nitrogen-fixing bacteria [7, 68]. In the process of transport, Sugiyama *et al*. [69] revealed that the soybean ABC transporters play important roles in legume-rhizobium symbiosis, and Clarke *et al*. [70] found by proteome analysis that transporters of sulfate, nitrate, peptides, and various metal ions like calcium, potassium and zinc are present on the soybean symbiosome membrane. Consistently, soybean ABC transporters and ion channels were predicted to interact with *B. diazoefficiens* proteins in the present study (Table 1). Since these transporters can facilitate the movement of nutrients between the symbionts and ensure the establishment of symbiosis, the candidate transport proteins in *G. max-B. diazoefficens* interactome can help our understanding of the role of transporters on the symbiosome membrane. In carbohydrate metabolism, soybean proteins involved in carbon metabolism, tricarboxylic acid cycle and N-glycan biosynthesis directly interacted with bacteria (Tables 2 and S7). Consistently, Libault *et al*. [68] and Carvalho *et al*. [7] showed that carbohydrate metabolism like the tricarboxylic acid cycle and glycolysis were induced by the presence of rhizobia in both roots and root hairs. These metabolic effects ensure the development of nodules by providing the carbon [71], while the host plant provides rhizobia with all the essential nutrients such as carbon required for bacterial metabolism [38]. Various signal transduction pathways play important roles in various stages of the symbiosis. They can coordinate the development of epidermal and cortical cells to ensure rhizobial invasion and nodule initiation [7, 72]. Previous studies have confirmed the involvement of many nod factors in the signal transduction processes such as G-protein coupled receptor signaling pathways [73, 74], small GTPase mediated signal transduction [75, 76], calmodulin [77], Soluble N-Ethylmaleimide Sensitive Factor Attachment Protein Receptor (SNARE) proteins [58, 78] and the MAPK (Mitogen-activated protein kinase) cascade [79]. In the present study, 34 G-proteins, 18 small GTPase, 32 calmodulin and 18 SNARE proteins were present in the predicted PPIs and directly interacted with bacteria (Tables 1 and S8). The subnetworks of related signaling transduction provide opportunities to reveal whether and how these networks are interconnected, and then give insights into the mechanism of symbiosis.

### Hubs in the predicted network played roles in symbiosis

In previous studies, HSPs were reported to be involved in the host-pathogen interaction [80] and to be induced during symbiosis in response to pathogens [81–83], suggesting that HSPs play critical roles in the response of plant cells to biotic stressors. HSPs have also been identified in the symbiosome membrane of soybean [84], *L. japonica* [63] and *M. truncatula* [85] by proteome analysis. Moreover, Brechenmacher *et al*. [6] reported that HSPs were up-regulated in soybean roots during the interaction between *G. max* and *Bradyrhizobium japonicum*. In the present study, five of the top ten soybean hubs interacting with *B. diazoefficens* are HSPs, and three hub HSPs interacted with *B. diazoefficens* proteins in the metabolism of glycerophospholipids, an important component of bacteria membrane lipids (Table S8). The results in this study give us insights that the five HSPs interacting with bacteria in the predicted PPIs are key players in the establishment of RNS.

The other two highly interacting hubs were SGF14k and SGF14g, which were shown to interact with *B. diazoefficens* proteins in the pathways of two-component systems (TCSs) and tryptophan metabolism (Table S10). TCSs are abundant signalling pathways in prokaryotes [86, 87]. They could transduce extracellular signals into the cell and regulate multiple cellular processes in response to environmental stimuli [88, 89]. More importantly, transcriptional regulators of TCS showed increased expression in bacteroids during RNS [39]. For Tryptophan metabolism, Hunter [90] showed that *Bradyrhizobia* with altered tryptophan metabolism frequently have altered symbiotic properties, and changes in the level of indole-3-acetic acid (a tryptophan metabolism product) that is involved in bacteria-plant interactions [6,91,92]. Notably, Radwan and Wu [62] revealed that two homologs of the above 14-3-3 proteins play critical roles in RNS. In the present study, subnetwork analysis showed that SGF14k and SGF14g interacted with four soybean nodulin genes (*Glyma.06G065600*, *Glyma.17G13300*, *Glyma.17G193800* and *Glyma.14G008200*). Among the four nodulin genes, two (*Glyma.06G065600* and *Glyma.17G193800*) were verified to interact with SGF14k and SGF14g by LCI assay experiments (Figures 3B and 4). Therefore, we deduce that *Glyma.14G176900* (SGF14k) and *Glyma.02G208700* (SGF14g) are involved in the process of nodulation.

Carbon metabolism was found to be closely related to RNS [7, 68]. Delmotte et al. [41] identified several proteins involved in carbon metabolism in symbiosome membrane of soybean, including a complete set of tricarboxylic acid cycle enzymes, gluconeogenesis and pentose phosphate pathway enzymes, by integrated proteomic and transcriptomic analysis. In the present study, seven hubs (BAC49080, BAC52411, BAC45833, BAC47677, BAC47750, BAC45992 and BAC46205) were involved in carbon metabolism, including N-Glycan biosynthesis, Pyruvate metabolism, Glycolysis and Citrate cycle (Table S10). Additionally, enriched KEGG pathways contained protein processing in endoplasmic reticulum, Glycosylphosphatidylinositol (GPI)-anchor biosynthesis, Pentose phosphate pathway and Proteasome (Table S10). Yuan *et al*. [4] found that genes involved in protein processing in endoplasmic reticulum were differentially expressed between different developmental periods of the soybean nodule. Roux *et al*. [93] revealed that genes involved in GPI-anchor biosynthesis and proteasome function were found to be preferentially expressed in plant nodules. Therefore, these ten hubs of *B. diazoefficens* may be important symbiotic effectors and play roles in symbiosis.

### Subnetwork analysis provide insight into the mechanism of root nodule symbiosis

Subnetwork analysis of SNAREs showed that SNAREs mainly interacted with membrane transporters or related proteins (Figure 3A). In detail, Glyma.10G008300 interacted with a cation efflux system protein (BAC50315), two ABC transporter permease proteins (BAC51159 and BAC49765), a cation-transporting ATPase (BAC52318) and an ammonium transporter (BAC45878). Five SNARE proteins (Glyma.04G072700, Glyma.10G149000, Glyma.07G042400, Glyma.03G029700 and Glyma.01G137300) interacted with BAC49080, a cation-transporting ATPase. Glyma.10G149000 and Glyma.13G307600 interacted with a Na^+^/H^+^ exchanger (BAC46205). Sokolovski *et al*. [94] proved that a plasma membrane SNARE protein in *Nicotiana benthamiana* guard cells could regulate Ca^2+^ channels and also possibly target other ion channels. The results indicated that SNAREs in the symbiosome membrane may play roles in regulating bacteria ion channels. Further analysis of the role of SNARE proteins will provide novel insights into RNS.

Through the subnetwork analysis, two 14-3-3 proteins, SGF14k and SGF14g, not only interacted with soybean nodulins but also were closely connected with two bacterial DctA proteins, BAC48988 and BAC49563 (Figure 3B). DctA was an important transporter for C4-dicarboxylic acids, which are the main form of carbon and energy sources from host plant to *rhizobium* [95]. Notably, DctA was reported to be essential for symbiotic nitrogen fixation in *Sinorhizobium meliloti*, as well as other rhizobia [78, 96]. The relationships between the above two 14-3-3 proteins and DctA proteins were further verified by LCI assay experiments (Figure 4). Taken together, the results indicated that 14-3-3 proteins SGF14g and SGF14k regulate rhizobium DctA.

Of course, the predicted results are still far from complete and may inevitably contain a lot of false positives, as the coverage and accuracy of predicted PPIs largely depend on the quality of interaction data sets and the ability to identify the orthologs from the model organisms. Even so, the predicted PPI networks have allowed us to have an insight into the overall picture of the PPI network between *G. max* and *B. diazoefficiens* USDA 110, which provide useful information to understand the molecular mechanism of the legume-rhizobium symbiosis.

## Materials and methods

### Datasets

A collection of 8317 protein sequences of *B. diazoefficiens* USDA 110 were downloaded from the Ensembl genomes database (ftp://ftp.ensemblgenomes.org/pub/bacteria/release-30/fasta/bacteria_0_collection/bra dyrhizobium_diazoefficiens_usda_110/pep/) [97]. Soybean whole genome sequences (*G. max Wm82.a2.v1*) were obtained from Phytozome V10.3 (http://genome.jgi.doe.gov/pages/dynamicOrganismDownload.jsf?organism=PhytozomeV10) [98]. For genes with multiple transcripts, the longest protein sequence was chosen [99]. As a result, 56044 protein sequences were obtained in *G. max*.

To conduct the interolog analysis, we utilized the PPI information of seven well-studied model organisms, namely *Arabidopsis thaliana*, *Caenorhabditis elegans*, *Drosophila melanogaster*, *Escherichia coli* K12, *Homo sapiens*, *Mus musculus* and *Saccharomyces cerevisiae*. Experimentally verified PPIs of the aforementioned seven organisms were obtained from the public protein-protein interaction databases: BioGrid, DIP, HPRD, IntAct, MINT and TAIR (Table S1). The ortholog information between the aforementioned seven organisms and *G. max* or *B. diazoefficiens* independently were obtained from InParanoid 8 [100].

To carry out the domain-based PPI prediction, we downloaded the interacting Pfam domain pairs from the database of protein domain interactions (DOMINE Version 2.0) [101], which contains a total of 26219 domain-domain interactions (DDI). To increase the accuracy of prediction, only 2989 high-confident domain pairs were used as reference in this study.

### PPI prediction

Our PPI prediction was mainly based on the interolog method, along with the domain-based method to improve prediction accuracy. In the interolog method proposed by Walhout *et al*. [25], the pair of interactions A–B and A1–B1 are called an interolog if interacting proteins A and B in a species have interacting orthologs A1 and B1 in another species. Based on this theory, interolog PPI prediction is a process that maps interactions in the source organism onto the target organism to find possible interactions [26]. In the domain-based method, the two proteins are expected to interact with each other if a protein pair contains at least one interacting domain pair [27]. The protein domain annotations for *B. diazoefficiens* USDA 110 were conducted in the Pfam website [102], and the annotations for *G. max* proteins were obtained from Phytozome V10.3.

In this study, ortholog pairs between each of the aforementioned seven model organisms and *G. max* (or *B. diazoefficiens*) were obtained from the InParanoid database [100]. InParanoid scores between 0 and 1 reflect the relative evolutionary distance between orthologous gene pairs [103, 104]. The top score 1.0 is the best blast hit and has high credibility, and orthologs with scores below 1.0 are more or less sensitive. To restrict the sensitivity, ortholog pairs were selected with a score cutoff of 0.5. These ortholog pairs were further divided into two groups according to InParanoid score: orthologs with top score 1.0 and ones with a score between 0.5 and 1.0. Using the interolog method, all the above ortholog pairs were mapped onto the integrated PPI interactomes of the seven model organisms to predict PPIs. The predicted PPIs with low confidence orthologs were further filtered by the domain-based method to increase prediction accuracy and to decrease false positives (Figure 1).

Identification of secreted and membrane proteins in *B. diazoefficiens* USDA 110

The transmembrane and secreted proteins in *B. diazoefficiens* USDA 110 are considered to be positive candidates for interactions with *G. max*. All the proteins in *B. diazoefficiens* USDA 110 were used to predict transmembrane proteins through TMHMM 2.0 [105] and to identify secretory proteins through SingleIP 4.0 [106]. In TMHMM 2.0, the proteins were inferred to be transmembrane if the number of predicted transmembrane helices was not <1, and the expected number of amino acids in at least one transmembrane helix was not <18. SingleIP 4.0 was employed with the default settings.

### GO annotation and measurement of functional similarity

The GO annotations of *B. diazoefficiens* USDA 110 and *G. max* were obtained from the Gene Ontology Annotation (UniProt-GOA) Database [107] and Phytozome V10.3 [98], respectively. Semantic similarity scores between GO terms were measured by Jiang and Conrath’s distance method and calculated in database FunSimMat [108, 109] to evaluate the reliability of the predicted PPIs [110]. Jiang and Conrath’s distance between two GO terms is based on information content and was defined as follows [111]:

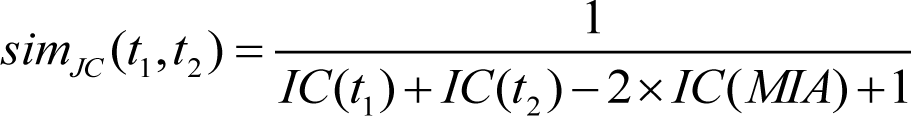

*sim_JC_*(*t*_1_, *t*_2_) is the set of common ancestors of terms *t*_1_ and *t*_2_ in the ontology, and ranges between 0, for no similarity, to 1, for highest similarity. We used sim_JC_ for referring to this score. As GO annotation classifies functions of a protein according to three features: molecular function, biological process and cellular component, there were, correspondingly, three independent sim_JC_ scores: sim_JC_^MF^, sim ^BP^ and sim ^CC^.

### Co-expression analysis

Transcriptome data of soybean were obtained from Phytozome V10.3 [98], which includes nine tissues (root, root hairs, nodules, leaves, stem, flower, pod, sam, and seed). The expression correlation between two interacting proteins was calculated using a widely used measure, Pearson correlation coefficient (PCC) [45]. The PCC value for each pair of non-self-interacting proteins was calculated using the Fragments Per Kilobase of transcript per Million mapped reads (FPKM) value of mRNA in the above nine tissues.

### Luciferase Complementation Image (LCI) assays for PPIs in *Nicotiana benthamiana* cells

#### Materials

Soybean (*G. max* Willimas 82) and tobacco plants were grown at 16-hlight / 8-h dark at 25°C for 30-60 d. *B. japonicum* (USDA110) was grown on (HM) medium plates at containing 50 μg of chloramphenicol/ml for selection of plasmid 25°C.

#### RNA and DNA Isolation

Soybean total RNA was isolated using the Trizol reagent (Invitrogen, Foster city, CA, USA) according to the manufacturer’s instructions and the RNAs were treated with the DNase I (Promega). The first-strand cDNA was then synthesized using M-MLV reverse transcriptase (Promega). The total DNAs of the *Bradyrhizobium japonicum* was isolated according to the method of Casse et al. [112].

#### Primers and conditions for PCR

Primers were analyzed by Oligo 6 (Table S11). PCR was carried out using a PCR system for 35 cycles (30 s at 95°C, 30 s at Tm and 1-4 mins at 72°C).

#### Luciferase Complementation Image (LCI) assays

Full length coding sequence of target genes were amplified by polymerase chain reaction from total RNA (Table S11) and were cloned into the *Bam*HI and *Sal*I sites of JW-771-N (NLUC), as well as *Kpn*I and *Sal*I sites of JW-772-C, to produce target gene-NLUC and target gene-CLUC recombination vectors for the LCI assay (for split Luc N-terminal/C-terminal fragment expression), respectively. Thus, N-gene, C-gene, N-LUC, and C-LUC were constructed according to previously described protocols. These constructs were transformed into Agrobacterium tumefaciens GV3101 strain through CaCl_2_ transformation [113]. The p19 protein (tomato bushy stunt virus) was used to suppress gene silencing [114].

#### Detection of interactions in vivo

The recombinant plasmids were transfected into *Agrobacterium tumefaciens* (GV3101). The OD600 of co-infiltrated *A. tumefaciens* strains is about 1.0 (gene-NLUC): 1.0 (gene-CLUC): 1.0 (P19), 500 μl of each, to co-culture for 2 h. Equal amount of the Agrobacterium suspension of each construct was mixed into a new 1.5 mL tube and vortexed for 10 sec to be ready for use. 8-10 weeks-old (16 h-light and 8 h-dark) *Nicotiana benthamiana* leaves were used to inject *A. tumefaciens* cocultures described above. Placed the tip end of the syringe (without needle) against the underside of the leaf (avoiding the veins) by supporting with one finger on the upperside, then gently pressed the syringe to infltrate the Agrobacterium mixture into the fresh leaf [115]. After growing for 48 h under the condition of 16 h-light and 8 h-dark, pieces of leaf abaxial epidermis were treated with 1 mM luciferin (promega, E1602), and the resulting luciferase signals were captured by Tanon-5200 image system (Tanon, Shanghai, China). To test each interacting protein pair, three experiments were performed and similar results were obtained.

## Supporting information

S1 Table. Experimental protein-protein interactions of seven model species from public databases

S2 Table. The selected 2,356 membrane and secreted proteins in *B. diazoefficiens* USDA 110

S3 Table. The predicted *G. max-B. diazoefficiens* interactome and detailed annotation information of the proteins, including 5115 inter-species PPIs between 2291 *G. max* and 290 *B. diazoefficiens* USDA 110 proteins

S4 Table. The predicted *G. max* interactome, including 233545 intra-species PPIs in soybean

S5 Table. The predicted *B. diazoefficiens* USDA 110 interactome, including 11106 intra-species PPIs in *B. diazoefficiens* USDA 110

S6 Table. List of 172 genes in the predicted PPIs that were detected to be expressed in bacteroids of the root nodule during symbiosis in at least one of three previous studies

S7 Table. List of input genes enriched in KEGG pathway enrichment analysis in Table 2 and their detailed annotation

S8 Table. Soybean proteins in the PPI network that were involved in signal transduction

S9 Table. Nodulation-related genes that experimentally interacted with *B. diazoefficiens* USDA 110 proteins

S10 Table. Top ten hubs of *G. max* and *B. diazoefficiens* USDA 110 in the *G. max-B. diazoefficiens* interactome and KEGG pathway enrichment analysis of these hubs by using their interacted proteins in the PPI interactome

S11 Table. Primers used in the luciferase complementation image (LCI) assays for PPIs in *Nicotiana benthamiana* cells

S1 Figure. Visualization of the predicted PPI network between soybean and *B. diazoefficiens* USDA 110. Each node represents a protein and each edge denotes an interaction. Red color circles represent soybean and yellow represent *B*. *diazoefficiens* USDA 110.

S2 Figure. Conserved PPIs identified in more than two species. Line represents the interaction relationship, circle represents proteins; yellow circles are *B. diazoefficiens* USDA 110 proteins, red, pink and grey circles are soybean proteins and respectively represent the expression values FPKM > 100, 5 < FPKM ≤ 100 and FPKM < 5 in nodules.

## Author contributions

YMZ conceived and designed the experiments, and revised the manuscript. ZXZ and PL assisted the supervision of the LCI experiment and bioinformatics analysis, respectively. LZ, HQ, ZBZ and MLZ performed bioinformatics analysis. JYL, YFD, JFZ performed the LCI experiments. YRC provided materials and modified the manuscript. LZ wrote the manuscript. All authors reviewed the manuscript.

