## Supplementary Materials for "*Bradyrhizobium diazoefficiens* USDA 110-*Glycine max* interactome provides candidate proteins associated with symbiosis"

1 **Supplementary Table S1**. Experimental protein-protein interactions of seven model species from public databases

| **Database/Species** | ***A. thaliana*** | ***C. elegans*** | ***D. melanogaster*** | ***E. coli K12*** | ***H. sapiens*** | ***M. musculus*** | ***S.cerevisiae*** |
|---|---|---|---|---|---|---|---|
| **BioGrid** | 35865(9100) | 8146(3879) | 42142(8116) | - | 203544(15598) | 13840(5431) | 260432(6010) |
| **DIP** | 394(316) | 4132(2712) | 23148(7622) | 12174(2867) | 4879(2980) | 1316(1191) | - |
| **HPRD** | - | - | - | - | 39205(9673) | - | - |
| **IntAct** | 15066(5656) | 17109(9635) | 41960(11296) | 17313(3186) | 97347(12590) | 18336(7741) | 85888(6125) |
| **MINT** | 240(222) | 5550(3293) | 22846(7386) | 23(29) | 20498(7610) | 2017(1557) | - |
| **TAIR** | 2177(1332) | - | - | - | - | - | - |
| **total** | **44702(9948)** | **28791(11543)** | **78383(9438)** | **24460(3358)** | **281387(15937)** | **31010(8567)** | **311333(6149)** |

2

**Supplementary Table S8.** Soybean proteins in the PPI network that were involved in signal transduction

| Protein classification | Genes |
|---|---|
| G-proteins | Glyma08g20610, Glyma04g02530, Glyma15g00930, Glyma15g40370, Glyma04g11560, Glyma20g37730, Glyma13g24140, Glyma01g03650, Glyma10g03660, Glyma06g16700, Glyma05g22480, Glyma13g44330, Glyma17g09980, Glyma09g04290, Glyma13g01270, Glyma19g28830, Glyma13g32940, Glyma04g05960, Glyma06g05960, Glyma14g11140, Glyma01g03650, Glyma01g43550, Glyma05g34120, Glyma10g29580, Glyma11g07330, Glyma02g01260, Glyma03g35470, Glyma06g02580, Glyma03g38600, Glyma04g01460, Glyma08g25673, Glyma15g40860, Glyma05g04210, Glyma06g08580, Glyma17g08100, Glyma03g28770, Glyma17g34450, Glyma15g11090, Glyma12g04810 |
| small GTPase | Glyma08g20610, Glyma04g02530, Glyma04g11560, Glyma20g37730, Glyma13g24140, Glyma01g03650, Glyma05g22480, Glyma17g09980, Glyma09g04290, Glyma13g01270, Glyma13g32940, Glyma01g03650, Glyma01g43550, Glyma10g29580, Glyma11g07330, Glyma02g01260, Glyma06g02580, Glyma06g08580, Glyma15g11090 |
| calmodulin | Glyma01g05440, Glyma02g11800, Glyma02g44350, Glyma03g00640, Glyma04g32470, Glyma05g13900, Glyma05g29050, Glyma05g38480, Glyma06g44510, Glyma08g12200, Glyma08g15150, Glyma08g16420, Glyma08g36780, Glyma08g38370, Glyma09g05110, Glyma09g38300, Glyma10g34430, Glyma10g36580, Glyma12g02850, Glyma12g13240, Glyma12g33280, Glyma13g27340, Glyma14g04460, Glyma14g37790, Glyma15g41650, Glyma15g42900, Glyma16g05101, Glyma16g19560, Glyma17g10270, Glyma19g19680, Glyma19g30140, Glyma20g35110 |
| SNARE proteins | Glyma04g07660, Glyma03g03490, Glyma17g35500, Glyma10g29020, Glyma16g08200, Glyma13g18030, Glyma18g38014, Glyma02g42280, Glyma02g35230, Glyma13g38370, Glyma13g19950, Glyma07g02870, Glyma01g33380, Glyma16g27050, Glyma07g04740, Glyma14g02120, Glyma13g33450, Glyma10g01131 |

**Supplementary Table S9.** Nodulation-related genes that interacted with *B. diazoefficiens* USDA 110 proteins

| **Soybean nod-related genes** | **Annotation of soybean genes** | ***B. diazoefficiens* USDA 110 genes** | **Annotation of *B. diazoefficiens* USDA 110 genes** | **Reference** |
|---|---|---|---|---|
| *Glyma.07G173700* | CHASE domain His kinase A | *BAC45924* | Two-component hybrid sensor and regulator | Murray *et al*., (2007)<br>Tirichine *et al*., (2007) |
| *Glyma.17G192100* | Early nodulin-like protein (ENOD93) | *BAC52652* | Hypothetical protein | Kouchi and Hata (1993) |
| *Glyma.04G060600* | Early nodulin-like protein (MtENOD20) | *BAC48391* | Cytochrome C-type biogenesis protein | Greene *et al*., (1998) |
| *Glyma.06G061200* | Early nodulin-like protein (MtENOD20) | *BAC48391* | Cytochrome C-type biogenesis protein | Greene *et al*., (1998) |
| *Glyma.14G101800* | Early nodulin-like protein (MtENOD20) | *BAC48391* | Cytochrome C-type biogenesis protein | Greene *et al*., (1998) |
| *Glyma.17G133000* | WD40 protein (MtCCS52) | *BAC46773* | HlyB/MsbA family ABC transporter | Vinardell *et al*., 2003 |
| *Glyma.08G120100* | Nodulin 26 | *BAC52381* | Aquaporin | Weaver *et al*., (1994) |
| *Glyma.04G250600* | Nodulin MtN21 | *BAC52381* | Aquaporin | Gamas *et al*., (1996) |
| *Glyma.13G091100* | Nodulin MtN21 | *BAC52381* | Aquaporin | Gamas *et al*., (1996) |
| *Glyma.U023000* | Nodulin MtN21 | *BAC52381* | Aquaporin | Gamas *et al*., (1996) |
| *Glyma.13G145100* | AtEIN2-like (MtSKL1) | *BAC50025* | Two-component hybrid sensor and regulator | Penmetsa *et al*., (2008) |
| *Glyma.06G065600* | Nodulin;SPFH/Band 7 family | *BAC46573* | Hypothetical protein | Winzer *et al*., (1999) |

**Supplementary Table S11**. Primers used in the luciferase complementation image (LCI) assays for PPIs in *Nicotiana benthamiana* cells

| **Gene** | **Primer** | **Sequence (5' to 3')** | **Gene** | **Primer** | **Sequence (5' to 3')** |
|---|---|---|---|---|---|
| *Glyma.17G193800* | 771-1F | CGAGCTCGGTACCCGGGATCCATGCCGTCGGACACCGTCGG | *Glyma.14G176900* | 771-10R | CGCGTACGAGATCTGGTCGACATTTTGTTCCCCGTCACCTT |
| *Glyma.17G193800* | 771-1R | CGCGTACGAGATCTGGTCGACTTCATCAAGTATGGCTTGAC | *BAC49735* | 771-AF | CGAGCTCGGTACCCGGGATCCATGTTCGACATCGCAGTCCC |
| *Glyma.06G065600* | 771-3F | CGAGCTCGGTACCCGGGATCCATGGAATTAAGCATAAAGAT | *BAC49735* | 771-AR | CGCGTACGAGATCTGGTCGACTAGATCGGCGGCGTTCGTTT |
| *Glyma.06G065600* | 771-3R | CGCGTACGAGATCTGGTCGACAGAACCATTATCGGGCAGGG | *BAC46573* | 771-BF | CGAGCTCGGTACCCGGGATCCATGTGCGGCACGCGCTCTTC |
| *Glyma.02G208700* | 771-5F | CGAGCTCGGTACCCGGGATCCATGGCAGCGAGCGCTCCCAC | *BAC46573* | 771-BR | CGCGTACGAGATCTGGTCGACCCCCTCCATCGCGTTCAGCC |
| *Glyma.02G208700* | 771-5R | CGCGTACGAGATCTGGTCGACATTTTGTTCCCCGTCACCTT | *BAC48988* | 771-DF | CGAGCTCGGTACCCGGGATCCGTGTCCGCAACAATCACTGC |
| *Glyma.13G158600* | 771-9F | CGAGCTCGGTACCCGGGATCCATGTGGCACGAGGCGAGGAG | *BAC48988* | 771-DR | CGCGTACGAGATCTGGTCGACGCCCGTCGTTACGGCGACCT |
| *Glyma.13G158600* | 771-9R | CGCGTACGAGATCTGGTCGACGTGCTTTGATCGATTTCTT | *BAC49563* | 771-EF | CGAGCTCGGTACCCGGGATCCATGACAACGACAACGATGGC |
| *Glyma.14G176900* | 771-10F | CGAGCTCGGTACCCGGGATCCATGCTTCCAAGTATGGATAA | *BAC49563* | 771-ER | CGCGTACGAGATCTGGTCGACGTCCGTCGTGATCGCCGTTT |

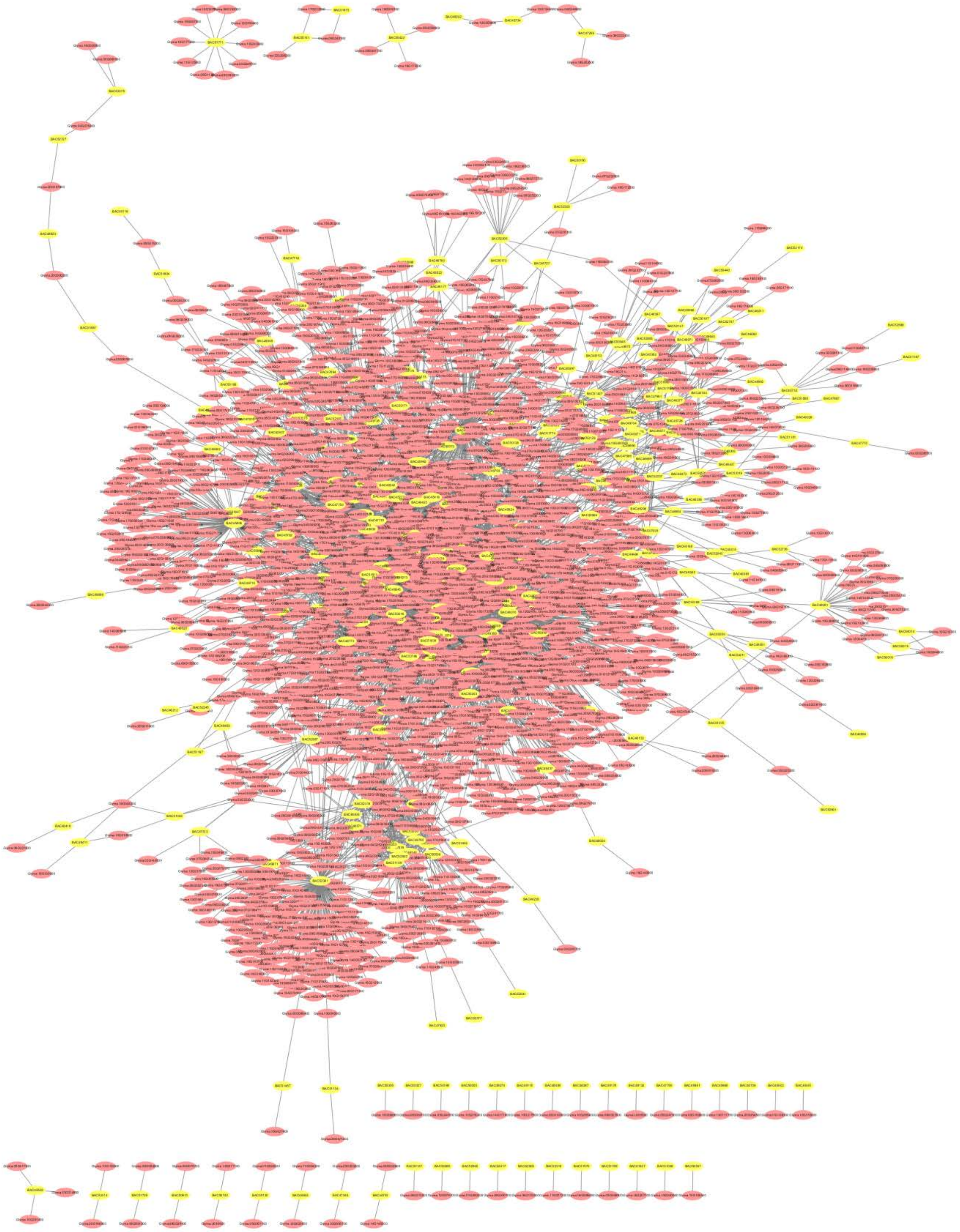

**Figure S1. Visualization of the predicted PPI network between soybean and *B. diazoefficiens* USDA 110**. Each node represents a protein and each edge denotes an interaction. Red color circles represent soybean and yellow represent B. *diazoefficiens* USDA 110.

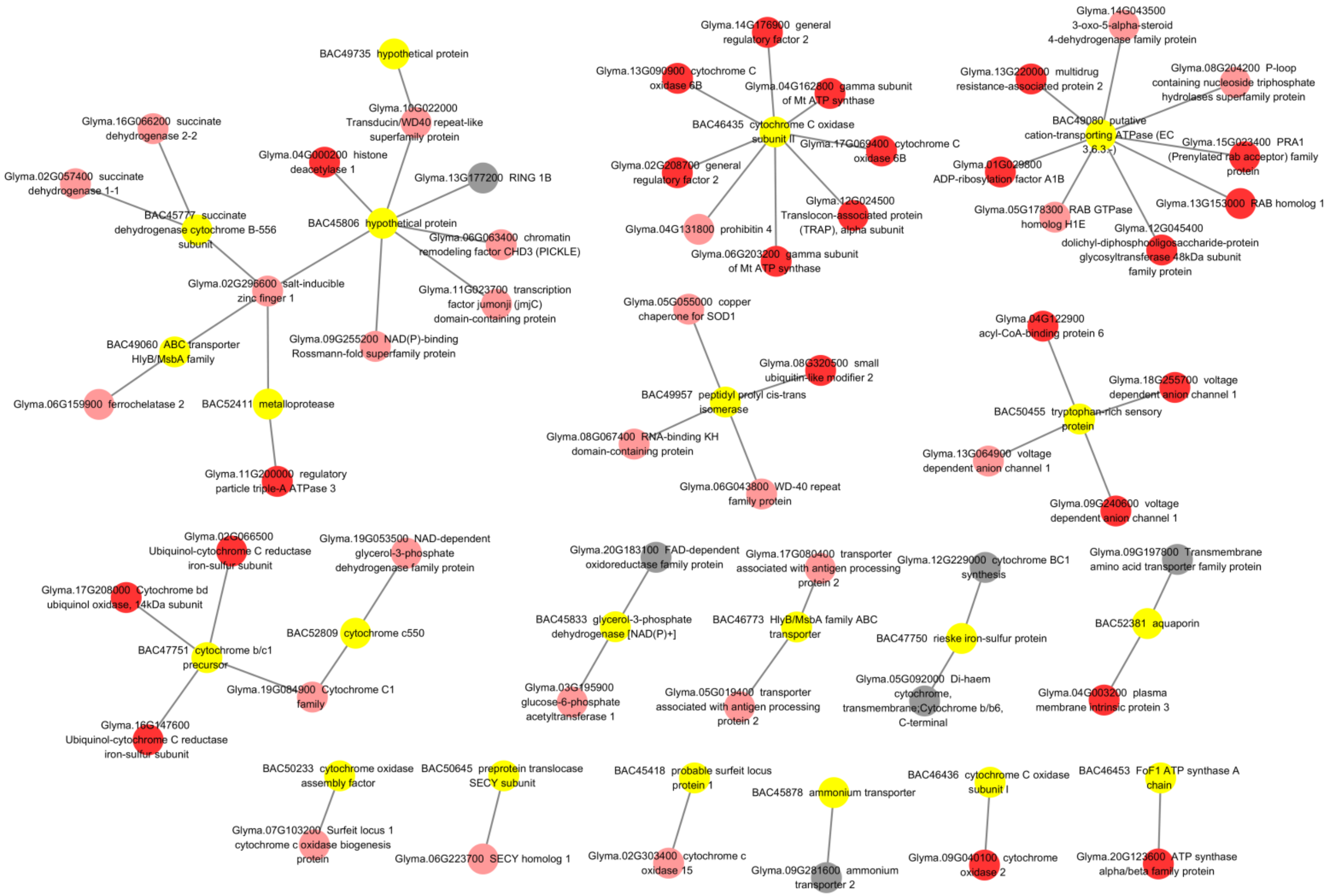


**Figure S2. Conserved PPIs identified in more than two species**. Line represents the interaction relationship, circle represents proteins; yellow circles are *B. diazoefficiens* USDA 110 proteins, red, pink and grey circles are soybean proteins and respectively represent the expression values FPKM > 100, 5 < FPKM ≤ 100 and FPKM < 5 in nodules.
